# Modulation of I-wave generating pathways with repetitive paired-pulse transcranial magnetic stimulation: A TMS-EEG study

**DOI:** 10.1101/2022.05.23.493173

**Authors:** Ryoki Sasaki, Brodie J. Hand, John G. Semmler, George M. Opie

## Abstract

**Objectives:** Repetitive paired-pulse transcranial magnetic stimulation (iTMS) at indirect (I) wave intervals increases motor-evoked potentials (MEPs) produced by TMS to primary motor cortex (M1). However, the effects of iTMS at early and late intervals on the plasticity of specific I-wave circuits remains unclear. The current study therefore aimed to assess how the timing of iTMS influences intracortical excitability within early and late I-wave circuits. To investigate the cortical effects of iTMS more directly, changes due to the intervention were also assessed using combined TMS-electroencephalography (EEG).

**Material and Methods:** Eighteen young adults (24.6 ± 4.2 years) participated in four sessions in which iTMS targeting early (1.5 ms interval; iTMS_1.5_) or late (4.0 ms interval; iTMS_4.0_) I-waves was applied over M1. Neuroplasticity was assessed using both posterior-to-anterior (PA) and anterior-to-posterior (AP) stimulus directions to record MEPs and TEPs before and after iTMS. SICF at inter-stimulus intervals of 1.5 and 4.0 ms was also used to index I-wave activity.

**Results:** MEP amplitude was increased after iTMS (*P* < 0.01) and this was greater for PA responses (*P* < 0.01), but not different between iTMS intervals (*P* = 0.9). Irrespective of iTMS interval and coil current, SICF was facilitated after the intervention (*P* < 0.01). While the N45 produced by AP stimulation was reduced by iTMS_1.5_ (*P* = 0.04), no other changes in TEP amplitude were observed.

**Conclusion:** The timing of iTMS failed to influence which I-wave circuits were potentiated by the intervention. In contrast, reductions in the N45 suggest that the neuroplastic effects of iTMS may include disinhibition of intracortical inhibitory processes.

## Introduction

Transcranial magnetic stimulation (TMS) is a non-invasive brain stimulation technique that is able to induce and measure neuroplastic changes in primary motor cortex (M1), providing important evidence for the flexibility of M1 neurons. Neuroplasticity involves alterations to glutamatergic and gamma-aminobutyric acid (GABA) neurotransmission (for review, see 1) and greatly facilitates physiological and functional recovery following brain injury, for example after stroke (for review, see 2) or traumatic brain injury (for review, see 3). Utilising TMS to modulate neuroplasticity after injury therefore has the potential to provide therapeutic benefits within neurorehabilitation.

When TMS is applied to M1, it produces a complex volley of waves within corticospinal neurons that summate at the spinal cord to produce a motor-evoked potential (MEP) (for review, see 4). The earliest component of this descending volley is the D-wave, which is thought to reflect direct activation of the corticospinal axon. This is followed by a series of I-waves that occur with a periodicity of ∼1.5 ms: these are referred to as early (I1) and late (I2 and I3) based on their recruitment order, and are thought to reflect input onto the corticospinal neuron from local interneuronal networks (5). While these waves can only be directly visualized using invasive recordings from the epidural space, it is possible to assess their activity using paired-pulse TMS. For example, when two stimuli are applied over M1 with an interstimulus interval (ISI) corresponding to the I-wave periodicity, the associated MEP is facilitated relative to the response generated by a single stimulus applied in isolation. This is referred to as short-interval intracortical facilitation (SICF) and is thought to index excitability of the I-wave circuits (for review, see 4).

While discrete application of paired-stimuli can index I-wave excitability, applying the same stimulus pairs repeatedly over a 15-minute period instead produces a robust increase in MEPs and SICF. This is referred to as I-wave periodicity repetitive TMS (iTMS) and is thought to induce long-term potentiation (LTP)-like changes in M1 (6-8). Interestingly, previous work has suggested that modifying the ISI used during iTMS can determine which I-wave circuits are influenced by the intervention (7). For example, short ISIs of 1.5 ms would influence the I1 wave circuitry, whereas longer ISIs of 4-5 ms would influence the I3 wave circuitry. As the early and late I-wave circuits have unique physiological and functional relevance (9, 10), an ability to target them selectively has important implications for the clinical application of brain stimulation interventions. However, the effects of iTMS timing on the activity of specific I-wave circuits has not been previously assessed.

The aim of the current research was therefore to investigate how iTMS applied with short and longer ISIs influences the excitability of early and late I-wave circuits. This was achieved by: *(1)* applying iTMS with ISIs of 1.5 ms (iTMS_1.5_, corresponding to the I1 wave) and 4 ms (iTMS_4.0_, corresponding to the I2-3 wave) in separate sessions and *(2)* measuring changes in MEPs and SICF using both posterior-to-anterior (PA) and anterior-to-posterior (AP) current directions, which are thought to recruit from different interneuronal populations (for review, see 11). As a secondary aim, we also sought to investigate the cortical response to iTMS more directly. This was achieved by using electroencephalography (EEG) to record the TMS-evoked EEG potential (TEP)(for review, see 12).

## Methods

### Participants

Eighteen healthy, young adults (7 men and 11 women; mean age ± SD = 24.6 ± 4.2 years; age range = 19-35 years) were recruited from the University and wider community to participate in this study. All participants were right-handed, free of neurological and psychiatric disorders, and were not taking any drugs that influence the central nervous system. Contraindications to TMS were assessed using the TMS adult safety screen (13). A nominal payment of $15 per hour was offered to compensate for time and cost of participation.

Written informed consent was provided prior to inclusion and this study was conducted in accordance with the *Declaration of Helsinki*. All experimental procedures were approved by the University of Adelaide Human Research Ethics Committee (approval number: H-026-2008).

### Experimental Arrangement

Each participant visited our laboratory for four experimental sessions that were approximately 2.5 hours long, held at the same time of day and separated by at least one week. Each session involved recording MEPs and TEPs before and after application of iTMS at either early or late intervals. While iTMS was always applied using a PA current, pre- and post-iTMS measures were recorded with PA and AP current in separate sessions (Figure 1). The order of the sessions was randomized within a participant. For the duration of each session, participants sat in a comfortable chair with their right hand pronated on a table and were instructed to keep their eyes open and remain relaxed. Surface electromyography (EMG) was recorded from the right first dorsal interosseous (FDI) muscle via disposable Ag/AgCl electrodes in a belly-tendon montage, with an additional Ag/AgCl electrode placed over the right ulnar styloid as an earth. EMG data were sampled at 2 kHz using a CED1401 interface (Cambridge Electronic Design, Cambridge, UK), amplified (1000×) and band-pass filtered (20–1000 Hz) by a CED1902 signal conditioner (Cambridge Electronic Design, Cambridge, UK). Line noise was removed using a Humbug mains eliminator (Quest Scientific, North Vancouver, Canada) and recordings were stored on a computer for off-line analysis.

**Figure 1.**
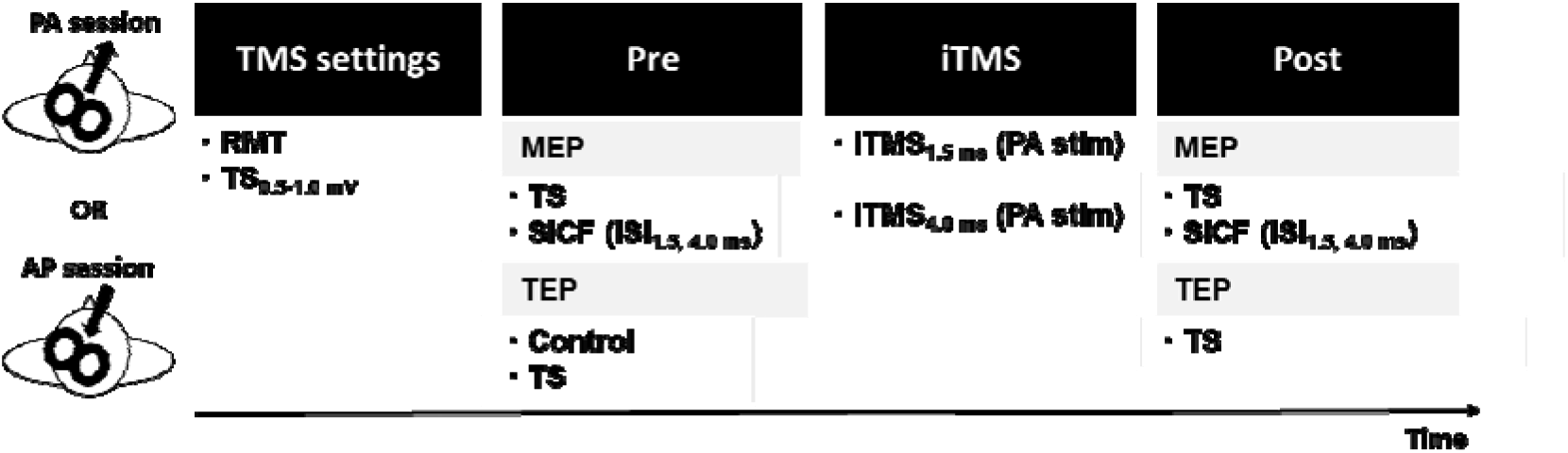
Experimental protocol. Four experimental sessions were performed involving iTMS sessions (iTMS_1.5_ and iTMS_4.0_) with a PA orientation and cortical assessments (both MEPs and TEPs) with PA and AP orientations separated by at least one week. Abbreviations; AP, anterior-posterior; ISI, inter-stimulus interval; iTMS, I-wave periodicity repetitive transcranial magnetic stimulation; MEP, motor-evoked potential; PA, posterior-anterior; SICF, short-interval intracortical facilitation; TEP, transcranial magnetic stimulation-evoked potential; TMS, transcranial magnetic stimulation; TS, test stimulus.

### TMS

Monophasic TMS pulses were delivered to the hand area of the left M1 using a figure-of-eight branding iron coil connected to two Magstim 200^2^ stimulators via a Bistim unit (Magstim, Dyfed, UK). The coil was held tangentially to the scalp at an angle of approximately 45° to the sagittal plane, at the location producing the largest stable response in the resting right FDI muscle. This position was co-registered to the MNI-ICBM152 brain template (14) using a Brainsight neuronavigation system (Rogue Research Inc, Montreal, Canada). Stimulation was applied at a rate of 0.2 Hz with a 10% jitter between trials. Resting motor threshold (RMT) was defined as the minimum intensity needed to evoke MEPs ≥ 50 µV in 5 of 10 consecutive trials during relaxation of the right FDI muscle (15). Stimulus intensity is expressed as a percentage of maximum stimulator output (MSO).

### SICF

SICF involved a subthreshold conditioning stimulus set at 90% RMT following a suprathreshold test stimulus (TS) at ISIs of 1.5 (SICF_1.5_) and 4.0 ms (SICF_4.0_), corresponding to the first and third SICF peaks (6, 16). The TS was set at the intensity required to produce an MEP of ∼ 0.5-1 mV when averaged over 20 trials. SICF at each time point was assessed using a single block of 60 trials (20 each of SICF_1.5_, SICF_4.0_, and TS), the order of which was pseudorandomised.

### iTMS

iTMS involved 180 pairs of stimuli applied in a PA orientation every 5 s, resulting in a total intervention time of 15 minutes (6, 8). The intensity was the same for both stimuli, and was adjusted so that paired stimulation produced a response amplitude of ∼1mV (assessed over 20 trials before the intervention). The ISIs targeting the first and third SICF peak (i.e., 1.5 and 4.0 ms) were applied in separate sessions (iTMS_1.5_ and iTMS_4.0_). In order to mitigate the effects of coil heating during the intervention, ice packs were used to cool the coil prior to and during iTMS application. This ensured that the same coil could be used for all TMS measures.

### EEG

EEG data was recorded using a WaveGuard EEG cap (ANT Neuro, Hengelo, The Netherlands), with 62 sintered Ag/AgCl electrodes in standard 10-10 positions, connected to an eego mylab amplifier (ANT Neuro, Hengelo, The Netherlands). CPz electrode was used as the reference for all recordings. Signals were filtered online (DC–0.26 × sampling frequency), digitized at 8 kHz, and stored on a computer for offline analysis. The impedance of all electrodes was constantly kept <10 kΩ through the experiment.

TEPs were recorded in a single block of stimulation that involved 100 pulses set at an intensity of 100% RMT, and this was always applied after measurement of MEPs. In an attempt to quantify the somatosensory- and auditory-evoked potentials that can confound the direct brain response, a block of shoulder stimulation was also recorded before iTMS (17, 18). This involved application of 100 TMS pulses set at 100% RMT, but with the coil held over the acromial process of the right shoulder. Although this approach cannot fully replicate the specific somatosensory input produced by TMS over the scalp, previous work has shown that the EEG response to shoulder stimulation accounts for much of the late TEP signal that is thought to be contaminated by somatosensory and auditory inputs (18), suggesting that this is an adequate control condition despite the different stimulation topography. In additional support of this approach, one recent study suggests that auditory input – which would have been comparable between scalp and shoulder stimulation in the current study – is the greatest source of sensory contamination to the TEP (19). During both scalp and shoulder stimulation, participants listened to white noise played through ear plugs to reduce the influence of auditory-evoked potentials. The volume of auditory masking was individually adjusted to minimize audition of the TMS click (18, 19).

### Data analysis

#### MEP data

MEP data were inspected visually and trials with muscle activity > 20 µV peak-to-peak amplitude in the 100 ms prior to TMS were rejected. MEP amplitude recorded in each trial was then quantified peak-to-peak and expressed in millivolts (mV). For SICF, the magnitude of facilitation recorded with each ISI was quantified as a percentage of the TS MEP amplitude recorded at baseline (8, 20). MEP amplitudes recorded during iTMS were averaged over 10 consecutive stimuli, resulting in a total of 18 blocks. All responses during iTMS were expressed relative to the mean response amplitude from the first block.

#### EEG data

All preprocessing and subsequent analysis was performed according to previously reported procedures (21, 22) using custom scripts on the MATLAB platform (R2019b, Mathworks, USA), in addition to EEGLAB (v2020.0) (23), TESA (v1.1.1.) (for review, see 22) and Fieldtrip (v20200607) (24) toolboxes. Data were epoched from -2000 ms to 2500 ms around the TMS trigger, baseline corrected from -500 ms to -5 ms and merged into a single file including both M1 (pre and post) and shoulder stimulation. Channels demonstrating persistent, large amplitude muscle activity or noise were manually removed, and then data segments associated with the large amplitude TMS artifacts were removed by cutting the data from -2 to 10 ms, and replacing it using cubic interpolation. The data was subsequently downsampled from 8 kHz to 500 Hz and epochs demonstrating bursts of muscle activity or electrode noise were semi-automatically removed. Interpolated data from -2 to 10 ms was then replaced with constant amplitude data (i.e., 0 s) and the conditions were split into two separate files (M1 and shoulder stimulation). An initial independent component analysis (ICA) was run on each condition using the FastICA algorism (25), and 1-2 independent components (IC’s) representing the tail of the TMS-evoked muscle artifact were removed (for review, see 22). Constant amplitude data from -2 to 10 ms were then replaced with cubic interpolation prior to the application of band-pass (1-100 Hz) and notch (48-52 Hz) filtering (zero-phase Butterworth filter implemented). In order to remove any additional decay artifacts still present after the first round of ICA, the source-estimate-utilizing noise-discarding (SOUND) algorithm was then applied; this approach estimates and removes artefactual components within source space, and also allows missing electrodes to be estimated and replaced (26). Following SOUND, data around the TMS pulse were again replaced with constant amplitude data prior to application of a second round of ICA, and IC’s associated with blinks, eye movements, electrode noise, and muscle activity were automatically identified using the TESA compselect function (default settings), and visually inspected prior to removal (for review, see 22). Data around the TMS pulse were then replaced with cubic interpolation, and all channels were re-referenced to average prior to a final baseline corrected (−500 ms to -5 ms).

### Statistical analysis

All analysis was performed using PASW statistics software version 28 (SPSS; IBM, Armonk, NY, USA) or Fieldtrip toolbox (EEG data only). Unless otherwise stated, data are displayed as mean ± SEM. Normality was assessed using Kolmogorov-Smirnov tests. Significance was set at *P* < 0.05.

### MEP data

Two-factor linear mixed model analysis with repeated measures (LMM_RM_) was used to compare baseline RMT, TS intensity, iTMS intensity, and TS MEP amplitudes between iTMS sessions (iTMS_1.5_ and iTMS_4.0_) and coil orientations (PA and AP). Three-factor LMM_RM_ was also used to compare baseline SICF between iTMS sessions, coil orientations and ISIs (SICF_1.5_ and SICF_4.0_). Two-factor LMM_RM_ was used to compare normalized MEP amplitudes during iTMS between iTMS sessions and blocks (B2-B18). For TS MEP amplitudes before and after iTMS, three-factor LMM_RM_ was used to compare values between iTMS sessions, coil orientations and time points (pre and post). Furthermore, four-factor LMM_RM_ was used to compare SICF between iTMS sessions, coil orientations, time points and ISIs. For all models, participant was included as a random effect, an AR(1) covariance structure was used, and restricted maximum likelihood estimation was applied. Each model also included single trial MEP data. Significant main effects and interactions were further investigated using custom contrasts with Bonferroni correction, implemented using the ‘Compare’ subcommand in SPSS.

### TEP data

In an attempt to identify the elements of the EEG signal that were likely to be more contaminated by sensory inputs, the TEP produced by M1 stimulation was compared to the response generated by shoulder stimulation in both spatial (i.e., between electrodes at each time point) and temporal (i.e., across time points within each electrode) domains using the Spearman correlation coefficient (17, 18). Spatial analyses were conducted from -50 to 350 ms, whereas temporal analyses were averaged over early (15-60 ms) middle (60-180 ms) and late (180-280 ms) time periods (17). For both measures, correlation coefficients were converted to Z-values using Fisher’s transform prior to group analysis (17, 19). Statistical significance was subsequently determined using a one-sample permutation test (derived from 10,000 permutations) assessing the hypothesis that each Z-score was greater than zero (i.e., positive correlation), with the t_max_ method used to control the family-wise error rate for multiple comparisons (27). The Z-values were transformed back into their original form for display (27). For data within each session, TEPs were compared between pre- and post-iTMS time points using cluster-based permutation analysis. Clusters were defined as two or more neighboring electrodes and 10,000 iterations were applied. A cluster was deemed significant if the cluster statistic exceeded *P* < 0.05 when compared with the permutation distribution. As correlation analysis demonstrated that TEPs were highly related to the response to shoulder stimulation from ∼60 ms post-TMS (see Fig 6), comparisons between conditions were limited to the early TEP components. This included N15 (10-15 ms), P30 (20-30 ms) and N45 (40-50 ms).

## Results

All 18 participants completed the sessions involving PA stimulation, but 3 participants had high stimulation thresholds that precluded collection of data with an AP orientation. Consequently, all measures for AP stimulation included data from 15 participants. No adverse events were reported. Baseline stimulus characteristics are compared between sessions and current directions in Table 1. Comparisons of RMT and TS intensity between coil orientations showed that stimulus intensities were all higher during AP stimulation (RMT: *F*_(1,26.17)_ = 82.98, *P* < 0.01; TS: *F*_(1,19.14)_ = 103.76, *P* < 0.01), but this was not different between iTMS sessions (RMT: *F*_(1,46.1)_ = 0.95, *P* = 0.34; TS: *F*_(1,45.83)_ = 1.65, *P* = 0.21) and there was no interaction between factors (RMT: *F*_(1,40.44)_ = 2.12, *P* = 0.15; TS: *F*_(1,38.30)_ = 0.39, *P* = 0.54). Baseline TS MEP amplitudes showed no differences between iTMS sessions (*F*_(1,432.52)_ = 0.001, *P* = 0.98) or coil orientations (*F*_(1,441.17)_ = 1.41, *P* = 0.24), and no interaction between factors (*F*_(1,480.74)_ = 0.29, *P* = 0.59). Comparisons of iTMS intensity showed higher intensities during the iTMS_4.0_ sessions (*F*_(1,18.656)_ = 5.35, *P* = 0.03) and during the PA sessions (*F*_(1,22.100)_ = 22.77, *P* < 0.01), but no interaction between factors (*F*_(1,42.173)_ = 0.59, *P* = 0.45). Comparisons of baseline SICF between ISIs showed that SICF_1.5_ resulted in greater facilitation than SICF_4.0_ (*F*_(1,570.74)_ = 260.36, *P* < 0.01). However, this was not different between iTMS sessions (*F*_(1,540.95)_ = 1.48, *P* = 0.22) or coil orientations (*F*_(1,543.02)_ = 0.77, *P* = 0.38), and there was no interaction between factors (all *P* > 0.18).

**Table 1.** TMS intensities and MEP amplitudes at baseline.

|  | iTMS<sub>1.5</sub> |  | iTMS<sub>4.0</sub> |  |
| --- | --- | --- | --- | --- |
|  | PA | AP | PA | AP |
| RMT (%MSO) | 56.6 ± 1.3 | 69.1 ± 1.7* | 57.1 ± 1.4 | 67.1 ± 1.8* |
| MEP<sub>0.5-1mV</sub> intensity (%MSO) | 67.0 ± 1.7 | 81.0 ± 1.7* | 66.5 ± 1.8 | 79.4 ± 2.0* |
| iTMS intensity (%MSO) | 57.2 ± 1.3 | 55.8 ± 1.3* | 62.1 ± 1.2<sup>†</sup> | 59.3 ± 1.3*<sup>†</sup> |
| MEP<sub>0.5-1mV</sub> amplitude (mV) | 0.74 ± 0.04 | 0.69 ± 0.04 | 0.71 ± 0.04 | 0.71 ± 0.04 |
| SICF<sub>1.5</sub> (%Test) | 254.9 ± 11.0 | 281.9 ± 14.6 | 304.7 ± 16.5 | 286.3 ± 13.0 |
| SICF<sub>4.0</sub> (%Test) | 124.1 ± 6.1<sup>#</sup> | 109.9 ± 6.9<sup>#</sup> | 117.0 ± 6.4<sup>#</sup> | 116.7 ± 6.8<sup>#</sup> |

\* $P < 0.05$ compared to PA stimulation. <sup>†</sup> $P < 0.05$ compared to iTMS<sub>1.5</sub>. <sup>#</sup> $P < 0.05$ compared to SICF<sub>1.5</sub>. Abbreviations; AP, anterior-posterior; iTMS, I-wave periodicity repetitive transcranial magnetic stimulation; MEP, motor-evoked potential; MSO, maximum stimulator output; PA, posterior-anterior; RMT, resting motor threshold; SICF, short-interval intracortical facilitation.

### Corticospinal excitability during iTMS

Figure 2 shows changes in MEP amplitude expressed as percentages relative to the first iTMS block. No difference was found between iTMS sessions (*F*_(1,2116.46)_ = 0.41, *P* = 0.52). However, values varied over blocks (*F*_(16,3207.57)_ = 4.87, *P* < 0.01), with post-hoc comparisons showing increased amplitudes during blocks 10-15, 17, and 18 relative to block 2 (all *P* < 0.03). Furthermore, there was an interaction between factors (*F*_(16,3204.25)_ = 1.90, *P* = 0.02), with post-hoc comparisons showing differences between iTMS_1.5_ and iTMS_4.0_ at B7, B12, B13, and B18 (all *P* < 0.05). Post-hoc comparisons also showed increased amplitudes during block 18 relative to block 2 in iTMS_1.5_ (*P* < 0.01) and during blocks 12-15 relative to block 2 in iTMS_4.0_ (all *P* < 0.02).

**Figure 2.**
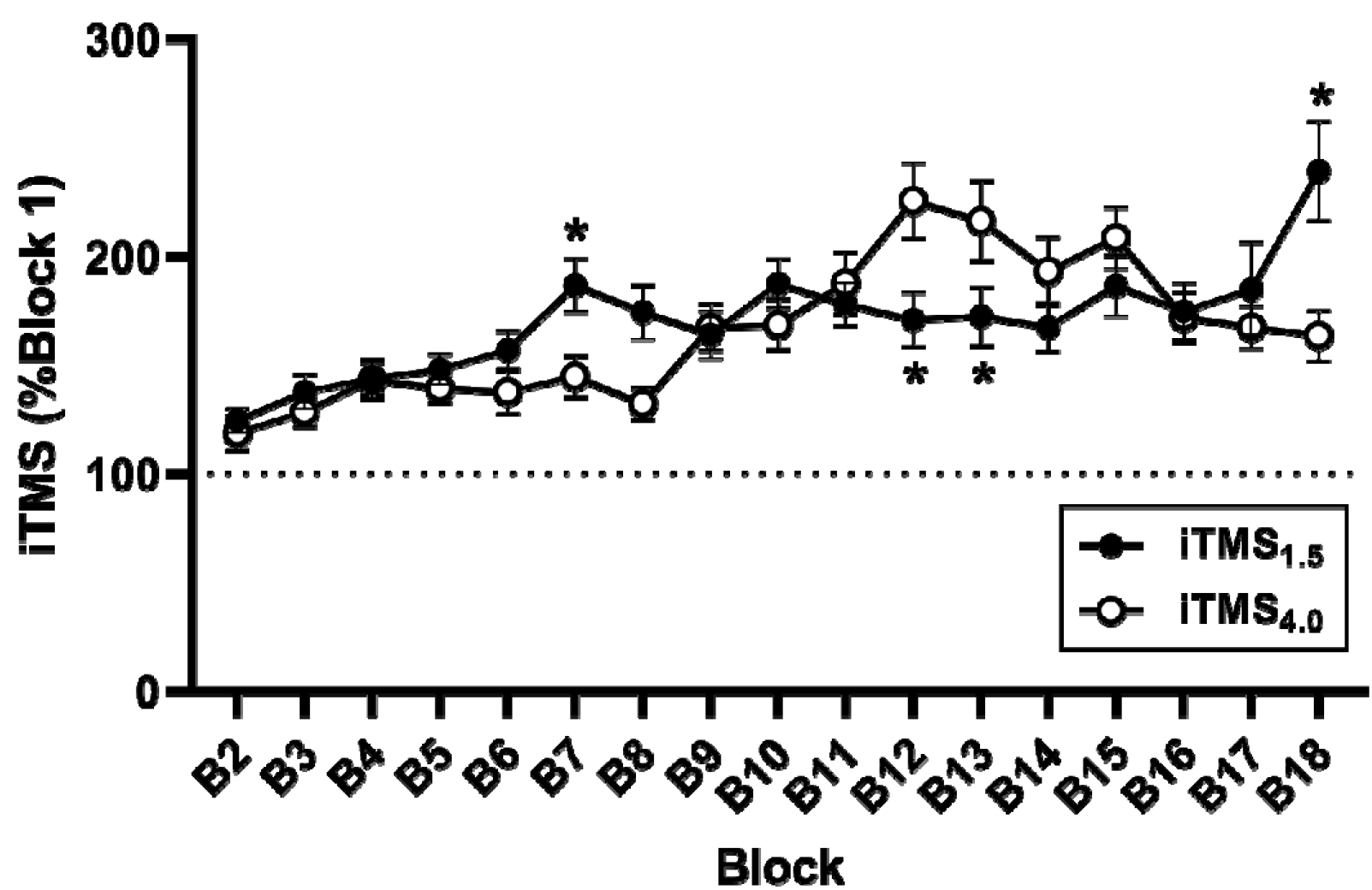
Corticospinal excitability changes during iTMS. iTMS_1.5_ (black circles) and iTMS_4.0_ (white circles) are averaged over 10 consecutive MEP trials, and then the block 2-18 were normalized by the first block. \**P* < 0.05 compared to iTMS_4.0_. Abbreviations; B, block; iTMS, repetitive paired-pulse TMS at I-wave intervals.

### Changes in corticospinal and intracortical excitability after iTMS

TS MEP amplitudes before and after iTMS are shown in Figure 3A and B. MEP amplitudes were not different between iTMS sessions (*F*_(1,635..68)_ = 0.02, *P* = 0.89). However, responses were larger with PA stimulation (*F*_(1,627.7)_ = 13.81, *P* < 0.01) and at the post-iTMS time point (*F*_(1,649.23)_ = 46.86, *P* < 0.01), and there was an interaction between coil orientation and time point (*F*_(1,642.07)_ = 4.16, *P* = 0.04). *Post-hoc* analysis showed that, although MEPs were increased after iTMS for both coil orientations (*P* < 0.01), post-iTMS responses were larger for PA than AP stimulation (*P* < 0.01). No other interactions between factors were found (all *P* > 0.44).

**Figure 3.**
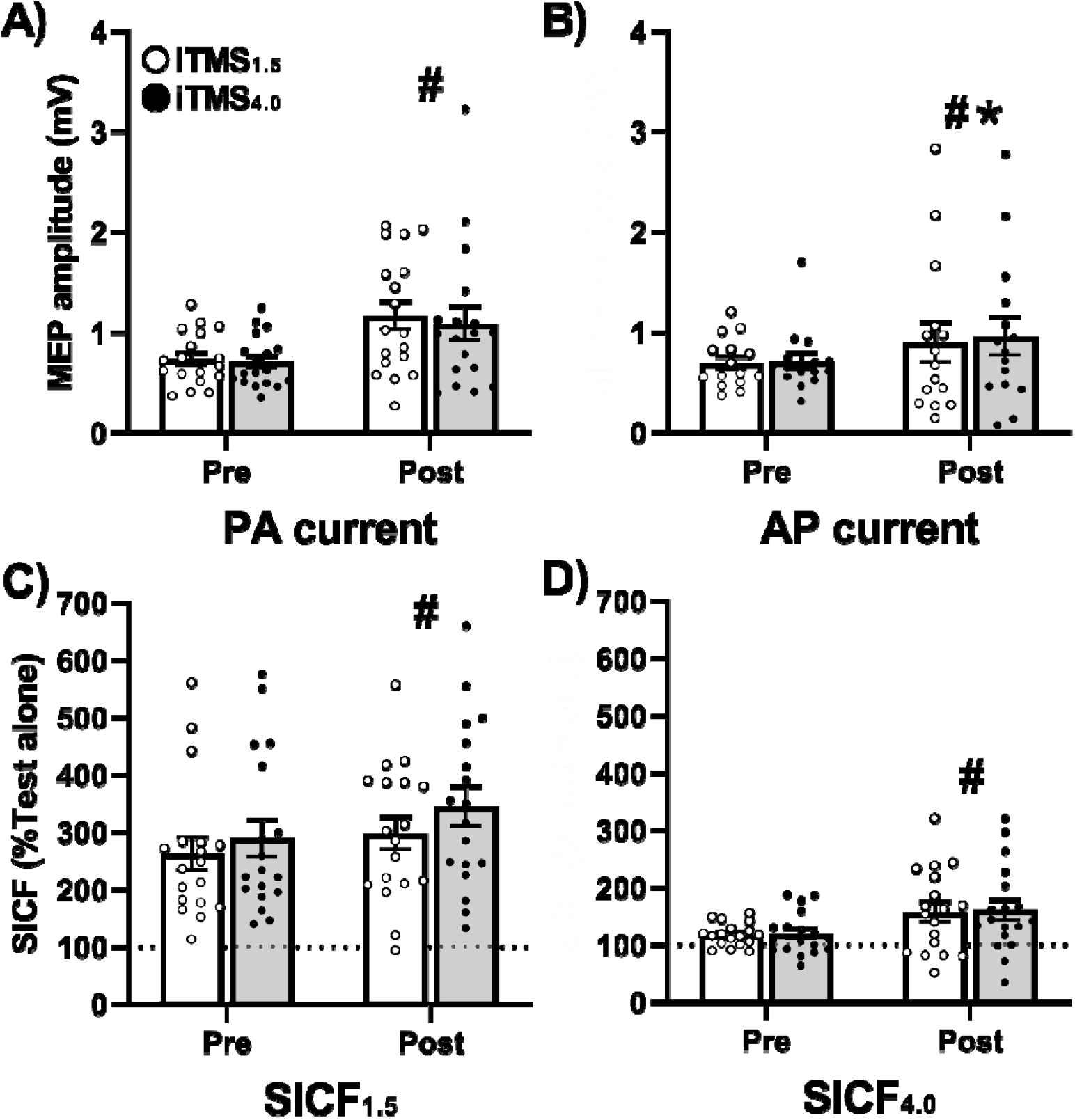
Corticospinal and intracortical excitability changes after iTMS. Top panels (*A, B*) represent TS MEPs with PA (*A*) and AP orientations (*B*) before and after iTMS_1.5_ and iTMS_4.0_. Bottom panels (*C, D*) represent SICF, which was normalized to baseline TS MEP amplitudes, with inter-stimulus intervals of 1.5 (*C*) and 4.0 ms (*D*) averaged over PA and AP orientations before and after iTMS_1.5_ and iTMS_4.0_. Each panel contains individual and mean values. ^#^*P* < 0.05 compared between pre and post; \**P* < 0.05 compared to PA responses at the same time point. Abbreviations; AP, anterior-posterior; iTMS, repetitive paired-pulse TMS at I-wave intervals; MEP, motor-evoked potential; PA, posterior-anterior; SICF, short-interval intracortical facilitation; stim, stimulation.

SICF before and after iTMS is shown in Figure 3C and D. While SICF was not different between coil orientations (*F*_(1,991.59)_ = 3.63, *P* = 0.06), it was increased after iTMS (*F*_(1,1017.89)_ = 27.3, *P* < 0.01), and varied between iTMS sessions (*F*_(1,989.45)_ = 7.5, *P* < 0.01) and ISIs (*F*_(1,1090.98)_ = 449.61, *P* < 0.01). Furthermore, there was an interaction between iTMS session and ISI (*F*_(1,1072.22)_ = 4.97, *P* = 0.03). Post-hoc analysis showed that SICF_1.5_ was larger than SICF_4.0_ within each iTMS session (*P* < 0.01), whereas SICF_1.5_ during the iTMS_4.0_ session was greater than during the iTMS_1.5_ session (*P* < 0.01). No other interactions between factors were found (all *P* > 0.13).

### TEPs preprocessing and correlation analysis

The average number of channels, epochs and IC’s removed during each step of the preprocessing pipeline are shown in Table 2. Figures 4 and 5 show grand-average TEP waveforms elicited by M1 and shoulder stimulation, whereas Figure 6 shows correlation coefficients resulting from comparisons between M1 and shoulder stimulation in both spatial (Figure 6A, B, C, D) and temporal (Figure 6E, F) domains. For both current directions, spatial correlations identified significant relationships between these conditions that began at ∼60 ms post TMS. In support of this, results of the temporal correlations suggested that the two signals were largely unrelated within the Early period, but became highly correlated across the scalp in Mid and Late periods. These results suggest that, although the early TEP response was likely to be less contaminated by sensory inputs, signal within the Mid and Late periods were likely to be heavily contaminated. Consequently, all statistical analyses of TEP amplitude were limited to the early period (Figure 7).

**Table 2.** Number of channels, epochs, and independent components removed during cleaning of TEPs.

|  | iTMS<sub>1.5</sub> |  | iTMS<sub>4.0</sub> |  |
| --- | --- | --- | --- | --- |
|  | PA | AP | PA | AP |
| Channels | $0.3 \pm 0.2$ | $0.4 \pm 0.2$ | $0.5 \pm 0.2$ | $0.3 \pm 0.2$ |
| Epochs (TS_pre) | $2.9 \pm 0.9$ | $5.6 \pm 2.0$ | $3.2 \pm 0.7$ | $5.5 \pm 1.5$ |
| Epochs (TS_post) | $4.4 \pm 1.2$ | $4.4 \pm 1.0$ | $4.6 \pm 0.9$ | $5.1 \pm 1.6$ |
| Epochs (Control) | $2.6 \pm 0.7$ | $2.3 \pm 0.5$ | $3.7 \pm 0.9$ | $4.0 \pm 1.6$ |
| ICA1 (TS) | $2.3 \pm 0.3$ | $2.5 \pm 0.5$ | $2.2 \pm 0.3$ | $2.0 \pm 0.2$ |
| ICA1 (Control) | $0 \pm 0$ | $0 \pm 0$ | $0 \pm 0$ | $0 \pm 0$ |
| ICA2 (TS) | $5.4 \pm 0.6$ | $6.1 \pm 0.6$ | $6.9 \pm 0.6$ | $5.5 \pm 0.7$ |
| ICA2 (Control) | $3.7 \pm 0.4$ | $3.1 \pm 0.3$ | $3.7 \pm 0.4$ | $3.1 \pm 0.3$ |

Abbreviations; AP, anterior-posterior; iTMS, repetitive paired-pulse TMS at I-wave intervals; IC, independent component analysis; PA, posterior-anterior; TS, test stimulus.

**Figure 4.**
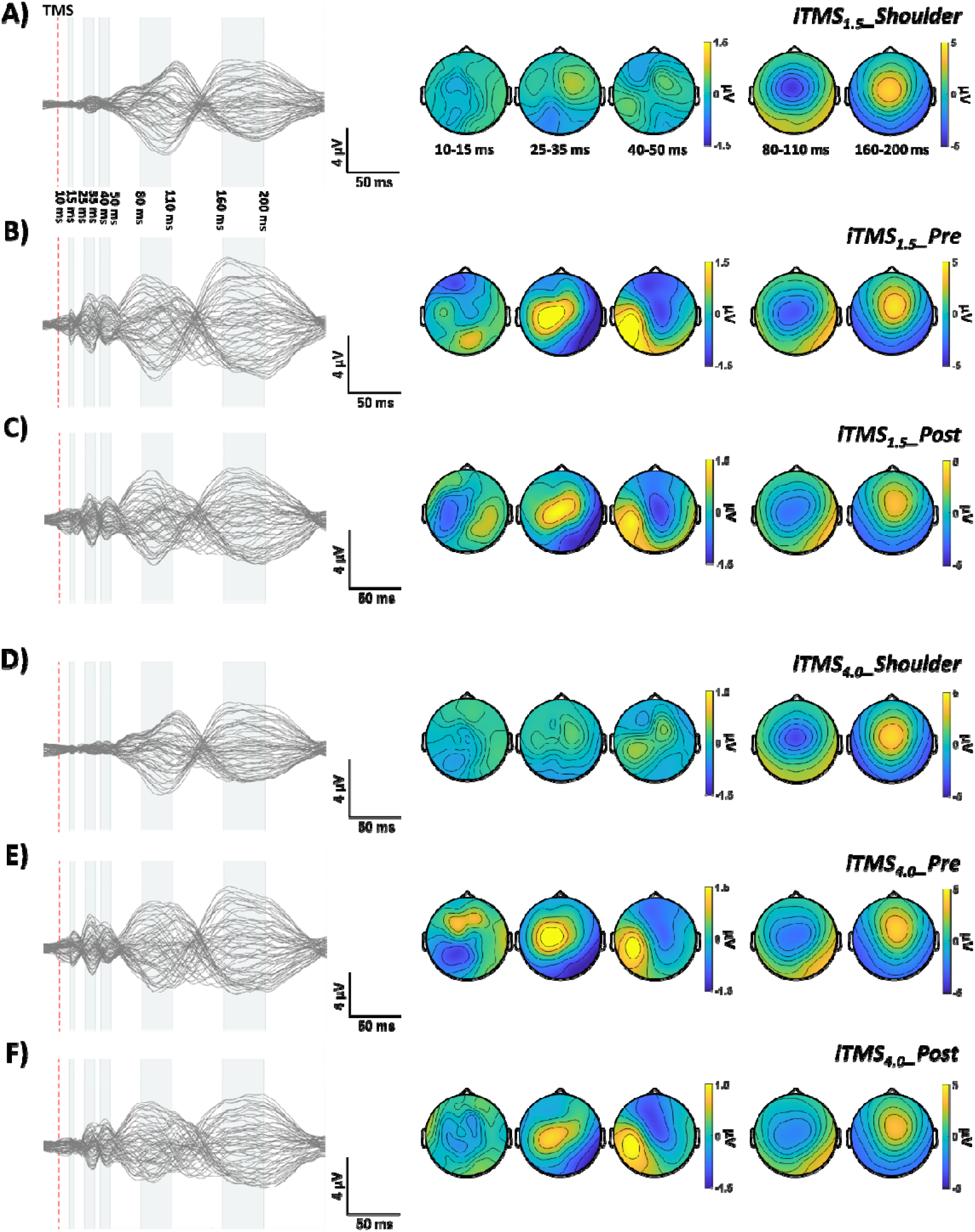
Grand average TEP waveforms and topographies with PA stimulation. (*A, B, C)* Shoulder (*A*) and M1 stimulation before and after iTMS_1.5_ (*B, C*). (*D, E, F)* Shoulder (*D*) and M1 stimulation before and after iTMS_4.0_ (*E, F*). Baseline TEP waveforms show several typical TEP components, named as N15, P30, P45, N100, and P180. Abbreviation; TMS, transcranial magnetic stimulation; iTMS, repetitive paired-pulse TMS at I-wave intervals.

**Figure 5.**
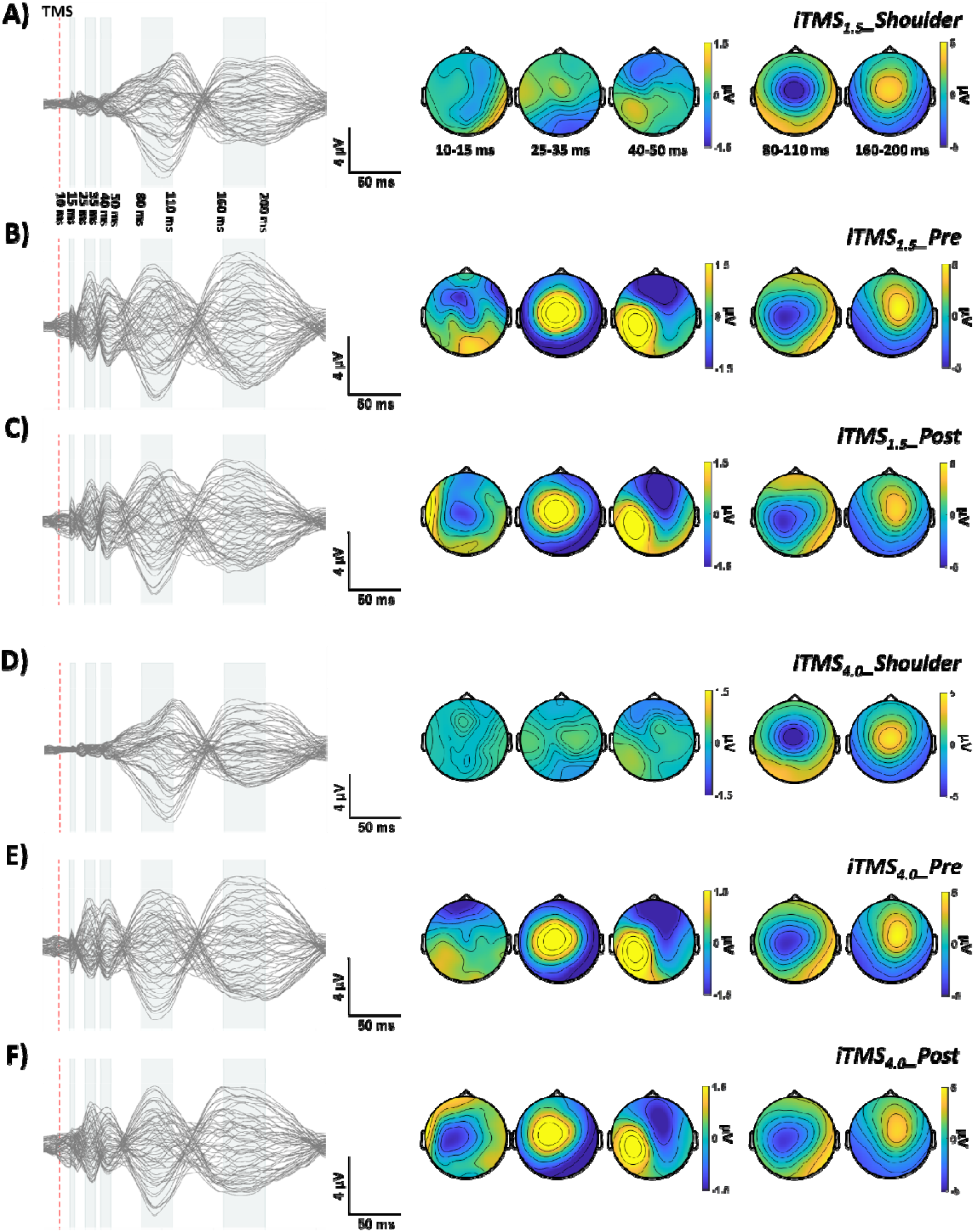
Grand average TEP waveforms and topographies with AP stimulation. (*A, B, C)* Shoulder (*A*) and M1 stimulation before and after iTMS_1.5_ (*B, C*). (*D, E, F)* Shoulder (*D*) and M1 stimulation before and after iTMS_4.0_ (*E, F*). Baseline TEP waveforms show several typical TEP components, named as N15, P30, P45, N100, and P180. Abbreviation; TMS, transcranial magnetic stimulation; iTMS, repetitive paired-pulse TMS at I-wave intervals.

**Figure 6.**
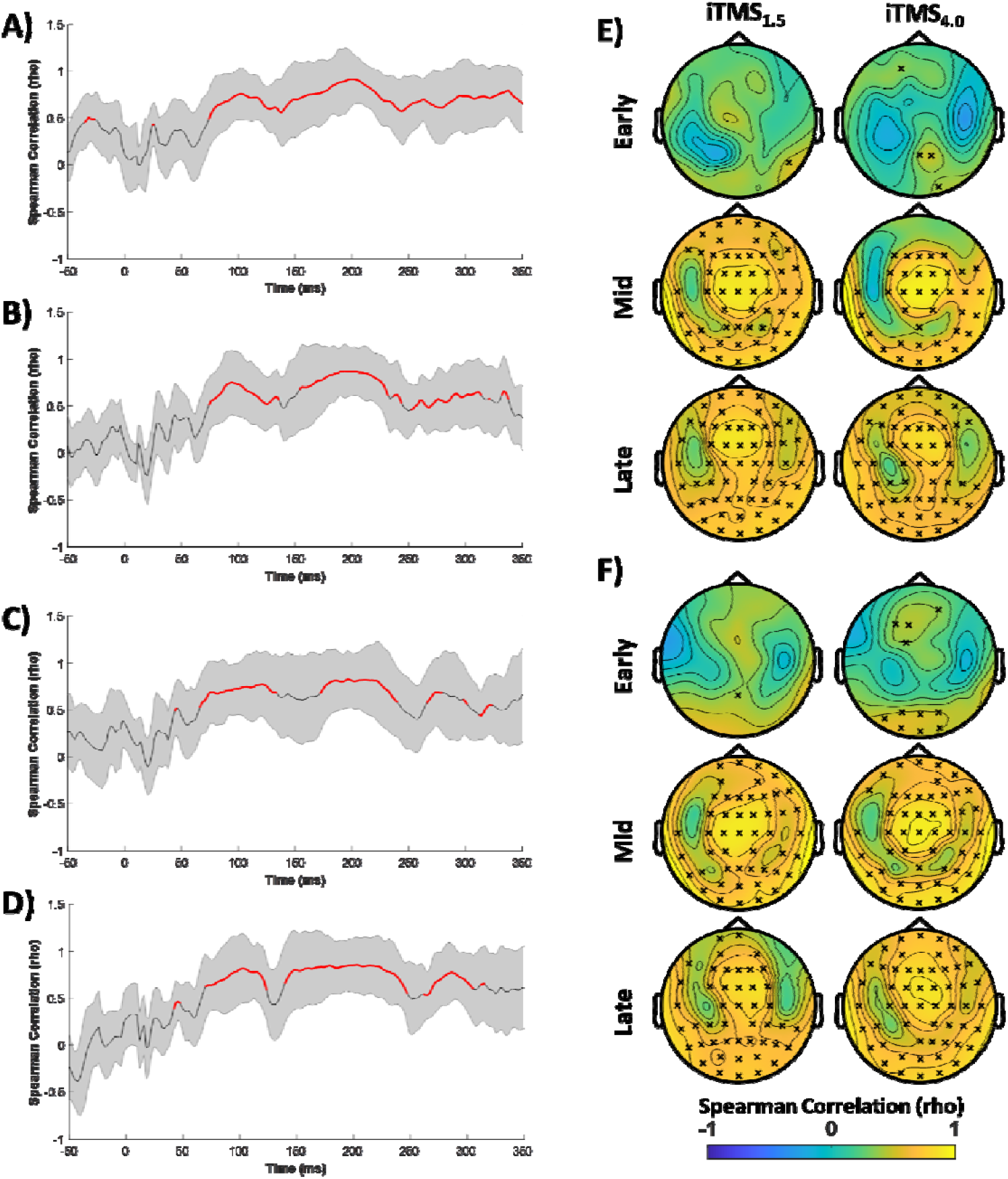
TEPs and sensory correlations. (*A, B, C, D*) Spatial correlations between EEG response to M1 and shoulder stimulation with PA current in iTMS_1.5_ *(A)* and iTMS_4.0_ *(B)* sessions and that with AP current in iTMS_1.5_ *(C)* and iTMS_4.0_ *(D)* sessions across all EEG electrodes. Red line segments indicate time periods that are significantly related between stimulation conditions. (*E, F*) Temporal correlations between EEG response to M1 and shoulder stimulation with PA (*E*) and AP (*F*) during Early (15-60 ms), Mid (60-180 ms) and Late (180-280 ms) time periods. Black crosses show that electrodes were significantly related between conditions. Abbreviation; iTMS, repetitive paired-pulse TMS at I-wave intervals.

**Figure 7.**
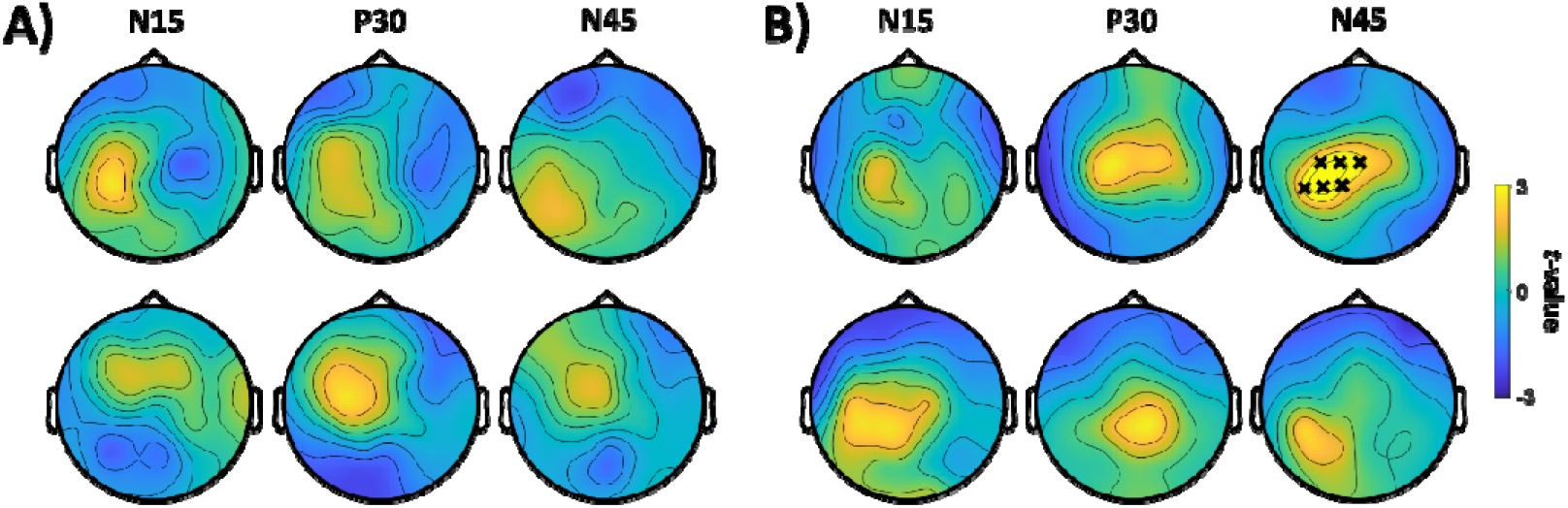
Comparison of TEPs between pre and post using cluster analysis. (*A, B*) Cluster-based permutation *t* test comparing the TEPs amplitudes with PA (*A*) and AP stimulation (*B*) before and after iTMS_1.5_ (top row) and iTMS_4.0_ (bottom row). Black crosses show a significant cluster between pre- and post-iTMS TEP amplitude.

### Changes in cortical excitability after iTMS

For PA sessions, there were no differences between pre- and post-iTMS TEP amplitude (all *P* > 0.06). In contrast, cluster-based comparisons of the N45 generated by AP stimulation identified a positive cluster (*P* = 0.039), which was associated with a decrease in amplitude after iTMS_1.5_. However, no differences were found for N15 and P30 (all *P* = 1). Furthermore, there was no change in any of the investigated components after iTMS_4.0_ (all *P* = 1).

## Discussion

The aim of this study was twofold: (1) to contrast the effects of iTMS applied with short and longer ISIs on the activity of early and late I-wave circuits and (2) to investigate the cortical response to iTMS. To achieve this, MEPs and TEPs were recorded using PA and AP current before and after iTMS_1.5_ and iTMS_4.0_. This approach produced facilitation of corticospinal (MEPs) and intracortical (SICF) excitability that was comparable between iTMS intervals. In contrast, changes in the TEP were only apparent after iTMS_1.5_, and were limited to the N45 produced by AP stimulation. While supporting the cortical effects of iTMS, these results also suggest that we were unable to specifically target different I-wave circuits by modifying the temporal profile of iTMS.

### Modifying iTMS ISI did not manipulate specific I-wave circuits

While previous work has investigated the effects of iTMS applied with short (20, 28) and longer (7, 8) ISIs, the current study is the first to compare these directly. In keeping with the existing literature, we found that iTMS with both intervals produced facilitation of MEPs and SICF, indicating a neuroplastic increase in M1 excitability. However, given that previous work has suggested that modifying ISI determines which I-waves are influenced by iTMS (7), we expected that the effects of iTMS would vary between ISIs. In particular, SICF is thought to provide a more specific index of excitability within different I-wave circuits (16), and we therefore expected its modulation by iTMS to be ISI-dependent (e.g., iTMS_1.5_ increases SICF_1.5_ but not SICF_4.0_, and vice versa). In contrast, changes to both MEPs and SICF were not different between iTMS ISIs. Consequently, our findings do not support the idea that modifying iTMS ISI allows specific targeting of different I-wave circuits.

While we were unable to demonstrate the expected specificity, it is important to note that stimulus intensities within the current study differed between SICF and iTMS. In contrast, previous work reporting differential effects of iTMS on specific I-waves used the same stimulus intensity for both. An alternative explanation for our results could therefore be that the neuronal populations targeted by our intervention may have differed to the population recruited by SICF, and this may have resulted in an apparent loss of specificity in how SICF was influenced by iTMS. In particular, di-synaptic disinhibition of an inhibitory circuit (likely involving gamma-aminobutyric acid type A; GABA_A_) has been shown to influence

I-wave excitability assessed with SICF at short and longer latencies (29). Furthermore, the perithreshold intensity we applied during iTMS would be expected to recruit relatively greater proportions of low threshold inhibitory circuits than the higher stimulus intensity used by Long and colleagues (7). Consequently, while the neuroplastic effects reported by Long and colleagues were likely more focused on the excitatory interneuronal circuits responsible for I-wave generation, it is possible that the effects of our intervention involved activation of both the low threshold disinhibitory circuit, and higher threshold excitatory circuits. Within this construct, activation of the disinhibitory circuit may have produced a generalised facilitation that obscured any temporally-specific effects of iTMS. While speculative, this possibility nonetheless demonstrates the importance of future work investigating the influence of stimulus intensity on the effects of iTMS.

In an attempt to more broadly characterise interneuronal circuits that might be differentially influenced by iTMS, excitability measures were recorded using both PA and AP currents. This approach found that single-pulse MEPs recruited with PA current were more potentiated than those recruited with AP current. One explanation for this response could be that the intervention was applied with a PA current, and elements activated by PA stimulation were therefore modulated to a greater extent. Given this, it remains possible that iTMS applied with an AP current many be more selective for modifying AP circuits. As this has not been attempted previously, it will be important to assess in future work. Nonetheless, the response within each current direction did not vary between iTMS intervals, further suggesting that modification to ISI did not influence specific I-wave circuits. A caveat to this interpretation is that stimulus conditions in the current study (i.e., 0.5-1 mV response in resting muscle) were unlikely to have produced isolated recruitment of early (PA current) or late (AP current) I-waves (for review, see 11). Consequently, our measures may not have been sensitive enough to identify subtle effects within different intracortical elements. Future work implementing more sensitive indices of I-wave recruitment (i.e., low intensity stimulation in active muscle) following iTMS will therefore be an interesting topic of investigation.

### TEP measures of cortical excitability are modulated by iTMS

Correlation analyses comparing TEP amplitude with the peripherally-evoked potential generated by shoulder stimulation suggested that responses were highly correlated from approximately 60 ms. This is consistent with a growing literature (17, 18), and has been suggested to indicate that the later TEP peaks are likely more contaminated by sensory-evoked potentials (17-19). To avoid the confounding influence of this contamination, we therefore decided to limit TEP analysis to the early components that are thought to be more reflective of cortical excitability, including N15, P30 and N45. The results of this approach suggested that the amplitude of N45 was reduced by iTMS (Fig 7). Studies using pharmacological intervention have suggested that N45 reflects activity of intracortical inhibitory circuits involving GABA_A_ (30-32). In support of discussion within the previous section, our TEP results therefore suggest that application of iTMS produced disinhibition of GABA_A_ergic inhibitory circuits.

As suggested above, the lower stimulus intensities we applied during iTMS may have resulted in effects on disinhibitory circuits that may not be as apparent following interventions applied with higher stimulus intensities. Consequently, it remains possible that utilising higher stimulus intensities during iTMS may reveal a different TEP response, possibly more focused on indices of motor cortical excitation like the P30 (for review, see 12). Despite this, it is interesting that changes in the N45 were only apparent in responses generated with AP stimulation following iTMS_1.5_. While the reason for this remains to be determined, it seems likely that it also reflects sensitivity to GABAergic circuits. For example, previous work using MEPs to assess short-interval intracortical inhibition (SICI) has shown that AP responses are more sensitive to activity of GABAergic inhibitory circuits, possibly due to preferential activation of late I-waves (17, 33, 34). Furthermore, the AP session of the iTMS_1.5_ intervention applied the lowest intensity stimulation (see *‘iTMS intensity’* in Table 1), suggesting that its activation of low threshold inhibitory circuits would have been relatively higher than the other sessions. Although speculative, this could suggest that manipulating stimulus intensity may be one way in which the effects of iTMS could be targeted to different intracortical circuits.

In conclusion, the application of iTMS with short and longer ISIs increased corticospinal and intracortical excitability, irrespective of iTMS interval. While these findings suggest that modifying the timing of iTMS has limited effects on which circuits are targeted by the intervention, clarification of how stimulus intensity influences contributions from intracortical inhibitory circuits is required. In support of this, iTMS also produced specific reductions in the N45 produced by AP stimulation, suggesting that disinhibition of GABA_A_ergic circuits contributes to the neuroplastic effects of this paradigm.

## Notes

**Source of financial support** RS is supported by Overseas Research Fellowship from the Japan Society for the Promotion of Science [grant number: 202060103]. GMO is supported by a National Health and Medical Research Council early career fellowship (APP1139723). Support was also provided by an Australian Research Council Discovery Projects Grant (grant number DP200101009).

**Conflicts of interest statement** None.

### Competing Interest Statement

The authors have declared no competing interest.

